# Selective targeting of type II tRNAs underlies SLFN14-mediated translational repression and its dysregulation by thrombocytopenia-linked mutations

**DOI:** 10.64898/2026.02.03.703619

**Authors:** Chengchao Ding, Xinyi Ashley Liu, Fushun Zhang, Saori Uematsu, Shu-Bing Qian, Yan Xiang

**Affiliations:** Department of Microbiology, Immunology and Molecular Genetics, Joe R. & Teresa Lozano Long School of Medicine, The University of Texas at San Antonio; 7703 Floyd Curl Drive, San Antonio, TX 78229, USA; Department of Infectious Disease, The First Affiliated Hospital of USTC, Division of Life Sciences and Medicine, University of Science and Technology of China; Hefei, Anhui, 230001, China; Division of Nutritional Sciences, Cornell University; Ithaca, NY 14853, USA

**Keywords:** tRNA, Schlafen, translation, endoribonuclease, thrombocytopenia

## Abstract

Schlafens (SLFNs) are interferon-inducible regulators of RNA metabolism and cell fate. SLFN14 is a ribosome-associated endoribonuclease whose pathogenic variants cause autosomal dominant inherited thrombocytopenia (IT), but the underlying disease mechanism has been unclear. We show that SLFN14 represses global protein synthesis through selective cleavage of type II tRNAs. IT-linked mutations alter RNA substrate specificity, enhancing degradation of type II tRNAs while reducing rRNA cleavage. This shift promotes ribosome stalling at codons decoded by type II tRNAs, triggering global translational arrest, stress signaling, and cell death. These findings define the molecular basis of SLFN14-associated thrombocytopenia and highlight selective tRNA targeting as a mechanism regulating translation and cell fate.

## Introduction

Schlafens (SLFNs) are a family of vertebrate genes with important roles in immune regulation, antiviral defense, and cellular sensitivity to DNA-targeted anticancer agents [1, 2]. Humans encode five SLFN proteins, SLFN5, SLFN11, SLFN12, SLFN13, and SLFN14. Among these, SLFN14 is unique for its association with inherited thrombocytopenia (IT). SLFN14 is predominantly expressed in hematopoietic lineages, especially megakaryocyte (MK) and erythroid precursors [3, 4]. Heterozygous missense mutations in SLFN14 cause an autosomal dominant form of IT, characterized by defective platelet function and excessive bleeding [5-8]. In addition to its role in hematopoiesis, SLFN14 has been identified as an interferon-stimulated antiviral factor that inhibits replication of influenza A virus [9], varicella zoster virus, and HIV-1 [10-12].

SLFNs are multidomain proteins that can be classified into subgroups based on domain architecture. Group III SLFNs, including SLFN5, SLFN11, SLFN13, and SLFN14, contain an N-terminal RNase domain, a central SWADL or linker region, and a C-terminal helicase-like domain, whereas group II SLFNs such as SLFN12 lack the helicase-like domain. Structures of several full-length SLFNs and isolated RNase domains have been determined [11-17], providing important insights into SLFN enzymatic activity and RNA substrate recognition. These studies revealed that the RNase domain adopts a U-shaped pseudodimeric architecture composed of N- and C-lobes, which together form a positively charged central valley capable of engaging RNA substrates. In vitro, RNase domains from multiple SLFNs display relatively broad and variable RNA substrate specificity. In particular, purified SLFN14 RNase domain cleaves rRNA, tRNA, and diverse structured RNAs containing double-stranded regions [11-13].

Notably, all known IT-associated SLFN14 mutations, including K218E, K219N/E, V220D, and R223W, cluster within the RNA-binding cleft of the RNase domain. Several of these mutations reduce RNA cleavage activity in vitro [11, 12]. For example, the K219N mutation reduces cleavage efficiency against both rRNA and tRNA [12]. However, despite extensive biochemical and structural characterization, the physiological RNA substrates through which SLFN14 exerts its cellular functions remain unclear. Moreover, the mechanisms by which IT-linked SLFN14 variants drive disease pathogenesis have not been resolved.

In this study, we show that SLFN14 represses global protein synthesis and identify selective cleavage of type II tRNAs as the primary mechanism underlying this activity. We further demonstrate that IT-linked mutations reprogram SLFN14 RNA substrate specificity, enhancing degradation of type II tRNAs while reducing rRNA cleavage. This altered RNA targeting promotes ribosome stalling, translational arrest, and cellular stress, thereby providing a mechanistic basis for SLFN14-associated thrombocytopenia.

## Results

### SLFN14 inhibits global protein synthesis, an effect exacerbated by IT-associated mutations

A consistent observation in SLFN14-linked IT is the dramatic reduction of SLFN14 expression in patient-derived platelets and in transfected cells [5-7]. This has been attributed to a dominant-negative effect of mutants on wild-type (WT) protein synthesis or stability. To broadly assess the effect of SLFN14 on translation, we transiently expressed SLFN14 in HEK293T cells and measured nascent protein synthesis using O-propargyl-puromycin (OPP) labeling. An mCherry tag was placed at the N-terminus of SLFN14 followed by a P2A ribosomal skipping sequence, allowing single-cell monitoring of SLFN14 expression by flow cytometry. Cells with no or low expression of WT SLFN14 showed protein synthesis comparable to untransfected controls (Fig. 1B). By contrast, cells with high levels of WT SLFN14 exhibited complete suppression of protein synthesis, similar to cycloheximide (CHX)-treated cells (Fig. 1B), indicating that SLFN14 overexpression potently inhibits translation.

**Figure 1.**
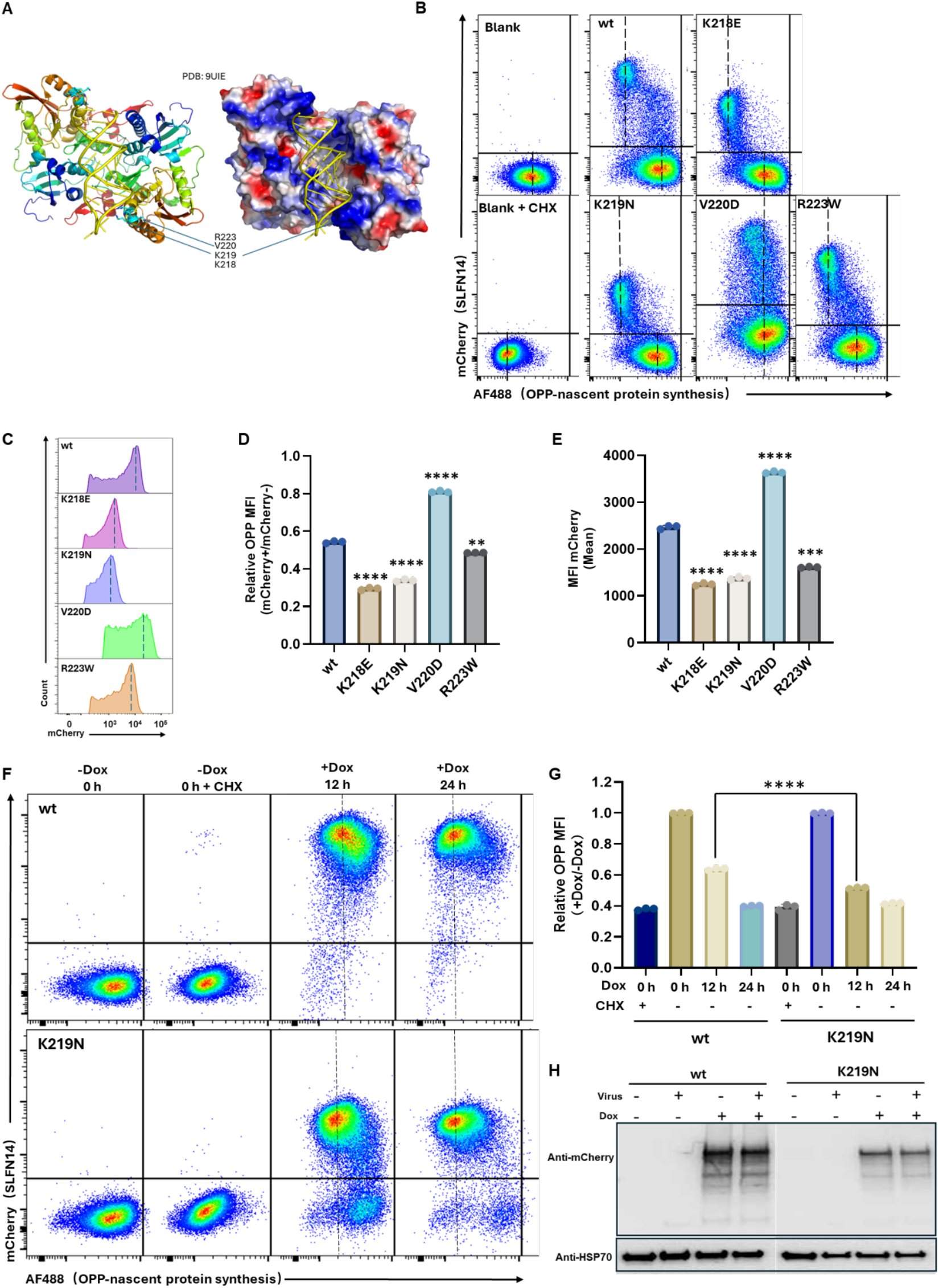
SLFN14 inhibits global protein synthesis, an effect exacerbated by IT-associated mutations. **(A)** Location of IT-associated mutations in the structure of the human SLFN14 RNase domain bound to dsRNA (PDB: 9UIE). The RNase domain is shown in both cartoon and surface representations, highlighting the positively charged central valley that accommodates the RNA substrate. Residues corresponding to IT-associated mutations are indicated. **(B-E)** HEK293T cells were transfected with mCherry-P2A-SLFN14 variants. At 16 h post-transfection, nascent protein synthesis was measured by OPP incorporation for 30 min. Untransfected cells and cells treated with CHX were included as controls. **(B)** Representative flow cytometry plots showing OPP levels relative to mCherry expression. The vertical dashed line indicates the mean fluorescence intensity (MFI) of the cell populations. **(C)** Histogram of mCherry-P2A–SLFN14 expression. **(D)** Quantification of nascent protein synthesis in SLFN14-expressing (mCherry^+^) cells relative to non-transfected (mCherry^-^) cells. **(E)** Quantification of mCherry MFI. **(F-H)** Dox-inducible 293T cell lines expressing mCherry-SLFN14 (WT or K219N) were left untreated or induced for the indicated times, and protein synthesis was measured as in (B). Representative plots are shown in **(F)**, with quantification of relative OPP MFI in Dox+ versus Dox-conditions in **(G). (H)** Western blot analysis of mCherry-SLFN14 expression in inducible cell lines using anti-mCherry and anti-HSP70 antibodies. “+ virus” indicates that the cells were infected with VACV. Statistical analysis was performed using one-way ANOVA (****P < 0.0001).

Strikingly, even cells expressing low levels of IT-linked variants K218E or K219N displayed little or no protein synthesis, demonstrating that these mutations enhance translational inhibition (Fig. 1B). The R223W variant also enhanced the translation inhibition, whereas the V220D variant reduced the inhibition. The extent of inhibition by the SLFN14 variants followed the order K218E ≈ K219N > R223W > WT > V220D (Fig. 1D). Notably, V220D has been reported to differ from other IT-linked mutations in that its expression did not reduce WT SLFN14 levels in transfected cells [5]. The effect of IT-linked mutations on protein synthesis was also reflected by mCherry expression levels in cells (Fig. 1C and E), as mCherry mean fluorescence intensity (MFI) correlated with OPP MFI.

To precisely control SLFN14 expression, we generated doxycycline (Dox)-inducible 293T cell lines expressing either WT or K219N mCherry-SLFN14. After 12 hours (h) of induction, protein synthesis was strongly reduced but not completely abolished, with the K219N variant showing greater inhibition than WT (Fig. 1F and G). After 24 h, both WT and K219N blocked protein synthesis, comparable to CHX treatment. Western blotting showed that the SLFN14^K219N^ protein level was substantially lower than WT (Fig. 1H), reflecting its stronger inhibition of cellular protein synthesis.

### SLFN14 inhibits vaccinia virus replication, with enhanced effects by IT-associated mutations

To test the effects of IT-associated mutations on antiviral activity, we examined their ability to restrict vaccinia virus (VACV), a cytoplasmic DNA virus. HEK293T cells were transfected with mCherry-P2A-SLFN14 constructs and infected with a GFP-expressing VACV. WT SLFN14 inhibited VACV replication, as indicated by reduced GFP expression in mCherry+ cells (Fig. 2A). The K218E, K219N, and R223W variants displayed even stronger antiviral activities, whereas V220D showed reduced antiviral activity compared to WT. The relative antiviral activities followed the order K218E ≈ K219N ≈ R223W > WT > V220D (Fig. 2B), which parallels their effects on global protein synthesis.

**Figure 2.**
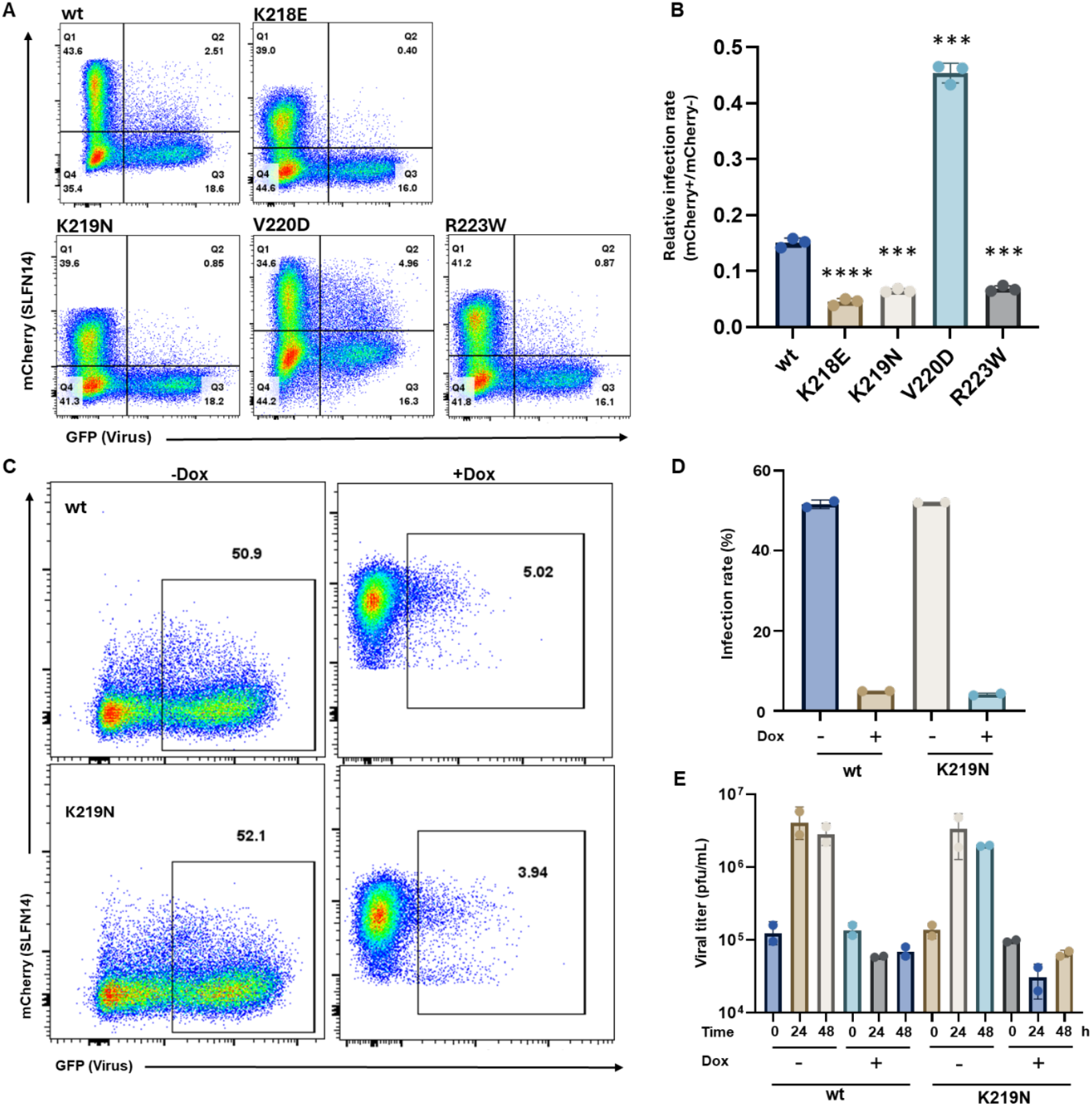
SLFN14 inhibits vaccinia virus replication, with enhanced effects by IT-associated mutations. **(A-B)** HEK293T cells were transfected with mCherry-P2A-SLFN14 variants for 36 h and infected with VACV/GFP^+^ for 15 h. **(A)** Representative flow cytometry plots showing GFP expression in SLFN14-expressing versus untransfected cells. Percentages of cells in each quadrant are indicated. **(B)** Relative infection (GFP^+^) rates between SLFN14-expressing (mCherry^+^) and nontransfected (mCherry^-^) cell populations were quantified from flow cytometry data. Rate = [Q2/(Q2 + Q1)]/ [Q3/(Q3 + Q4)]. **(C-D)** Dox-inducible 293T cell lines expressing mCherry-SLFN14 (WT or K219N) were left untreated or induced with Dox and subsequently infected with VACV/GFP^+^. Representative flow cytometry plots **(C)** and quantification of infection rates (GFP^+^ cells, 1= 100%) **(D). (E)** Viral titers at 0, 24, and 48 h post infection were measured by plaque assay on Vero cells. Statistical analysis was performed using one-way ANOVA (***P < 0.001, ****P < 0.0001).

We further examined antiviral activity against VACV with the Dox-inducible SLFN14 cell lines. Upon Dox induction, both WT and K219N lines exhibited a significant decrease in infection rates (%GFP+ cells) compared to uninduced controls (Fig. 2C&D). Viral titers measured by plaque assay confirmed that viral growth was completely abolished, with K219N cells showing a slightly greater reduction in viral titers than the WT cells at 24 h post infection (Fig. 2E).

### IT-associated mutations enhance SLFN14-mediated depletion of type II tRNAs while reducing rRNA cleavage

To investigate how IT-linked mutations alter SLFN14-mediated RNA processing, we analyzed total RNA from Dox-inducible WT and K219N cells. After induction for 12 hours, rRNA degradation was not detectable in either WT or K219N cells by nucleic acid staining (Fig. 3A). Surprisingly, type II tRNAs, with the longer variable loop, were found to be completely degraded in K219N cells while only slightly reduced in WT cells (Fig. 3B). This was confirmed by Northern blotting against the specific type II tRNAs (tRNA-Leu^CAA^, tRNA-Leu^TAA^, and tRNA-Ser^GCT^). The Northern blot also revealed that tRNA-Phe, a type I tRNA, was modestly reduced in K219N cells but remained unchanged in WT cells (Fig. 3C).

**Figure 3.**
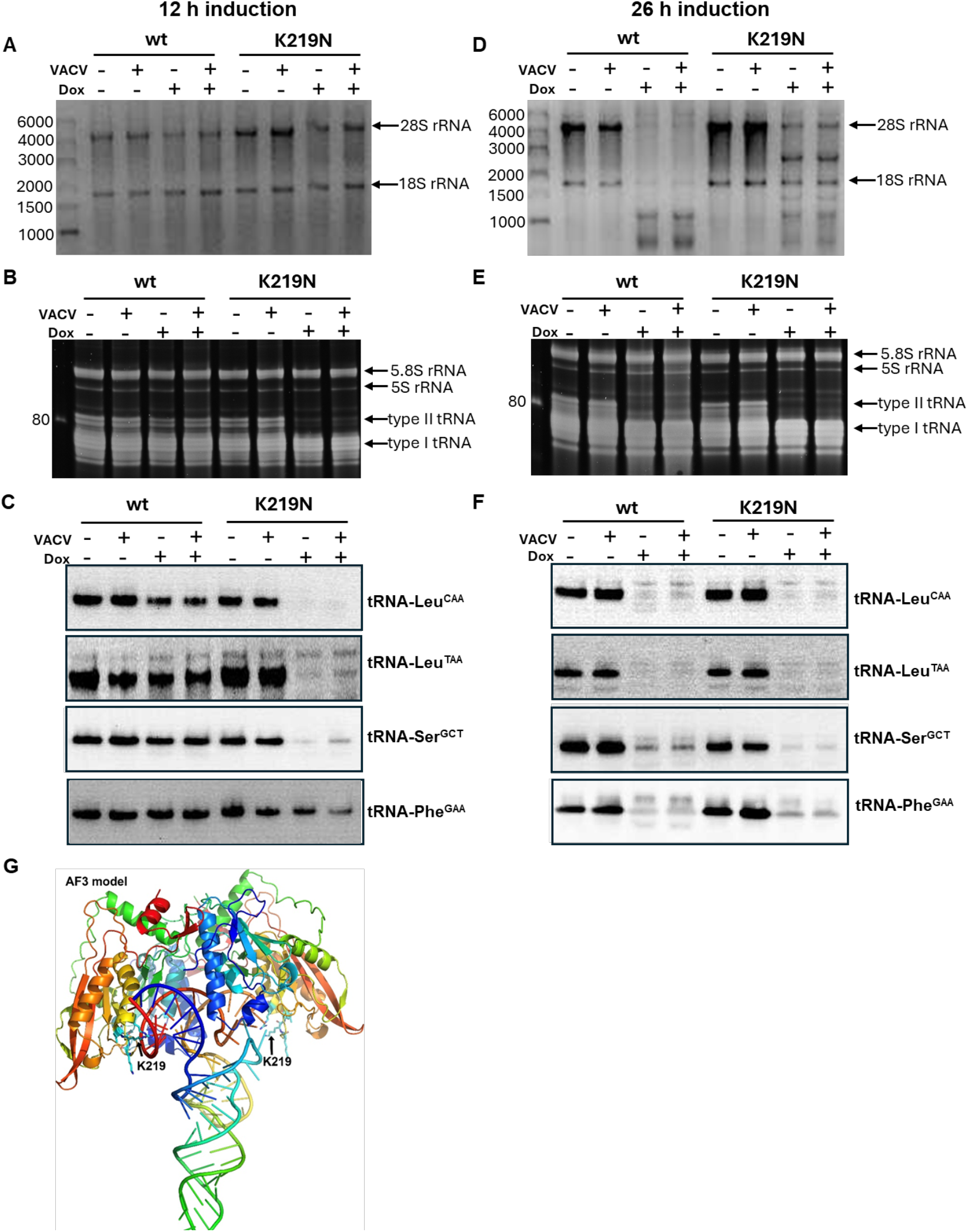
IT-linked mutations enhance SLFN14-mediated depletion of type II tRNAs while reducing rRNA cleavage. Analysis of total RNA from Dox-inducible 293T cells expressing WT or K219N SLFN14 after 12 h **(A-C)** or 26 h **(D-F)** of induction. **(A**,**D)** RNA was resolved on 2% agarose gel and stained with SYBR Safe DNA Gel Stain. RNA molecular weight markers (in nt) are shown. **(B**,**E)** RNA was resolved on a polyacrylamide/urea gel and stained with SYBR Gold stain. **(C**,**F)** Northern blot analysis using probes specific for the indicated tRNAs. VACV indicates infection with vK1^-^ C7^-^/GFP^+^ VACV. **(G)** AlphaFold3 model of SLFN14 RNase in complex with tRNA-Leu^TAA^. Residue K219 is highlighted and positioned in close proximity to the bound tRNA.

After induction of SLFN14 for 26 hours, rRNA degradation was prominent in WT cells but much weaker in K219N cells (Fig. 3D). Type II tRNAs were also greatly reduced in WT cells, although their levels remained higher than in K219N cells (Fig. 3E). While SYBR Gold staining could not clearly resolve changes in overall level of type I tRNAs, Northern blot demonstrated that tRNA-Phe was strongly reduced at this time point (Fig. 3F). In these experiments, an additional set of cells were also infected with VACV, and the results showed that the viral infection did not alter the RNA degradation by SLFN14.

Together, these results indicate that while SLFN14 targets both rRNA and tRNAs in cells, the K219N variant shifts substrate preference toward type II tRNAs. Consistent with this interpretation, an AlphaFold3 model of SLFN14 in complex with tRNA-Leu^TAA^ position K219 in close proximity to the bound tRNA (Fig. 3G), providing a structural explanation for the altered substrate specificity observed in the K219N mutant.

### SLFN14 causes ribosomal stalling at codons decoded by type II tRNAs, an effect exacerbated by IT-linked mutations

To assess the impact of SLFN14 on translation at codon resolution, we performed ribosome profiling (ribo-seq). Induction of WT SLFN14 for 24 hour caused ribosome stalling only at leucine codons TTA and TTG. In comparison, the K219N variant caused a stronger stalling at these codons, as well as stalling at leucine codons CTA and CTG and serine codons AGC and AGT (Fig. 4). Both serine codons are decoded by tRNA-Ser^GCT^. The pattern and magnitude of ribosome stalling correlated with the extent of tRNA degradation in WT and K219N cells, confirming that SLFN14 promotes ribosome stalling through preferential depletion of type II tRNAs.

**Figure 4.**
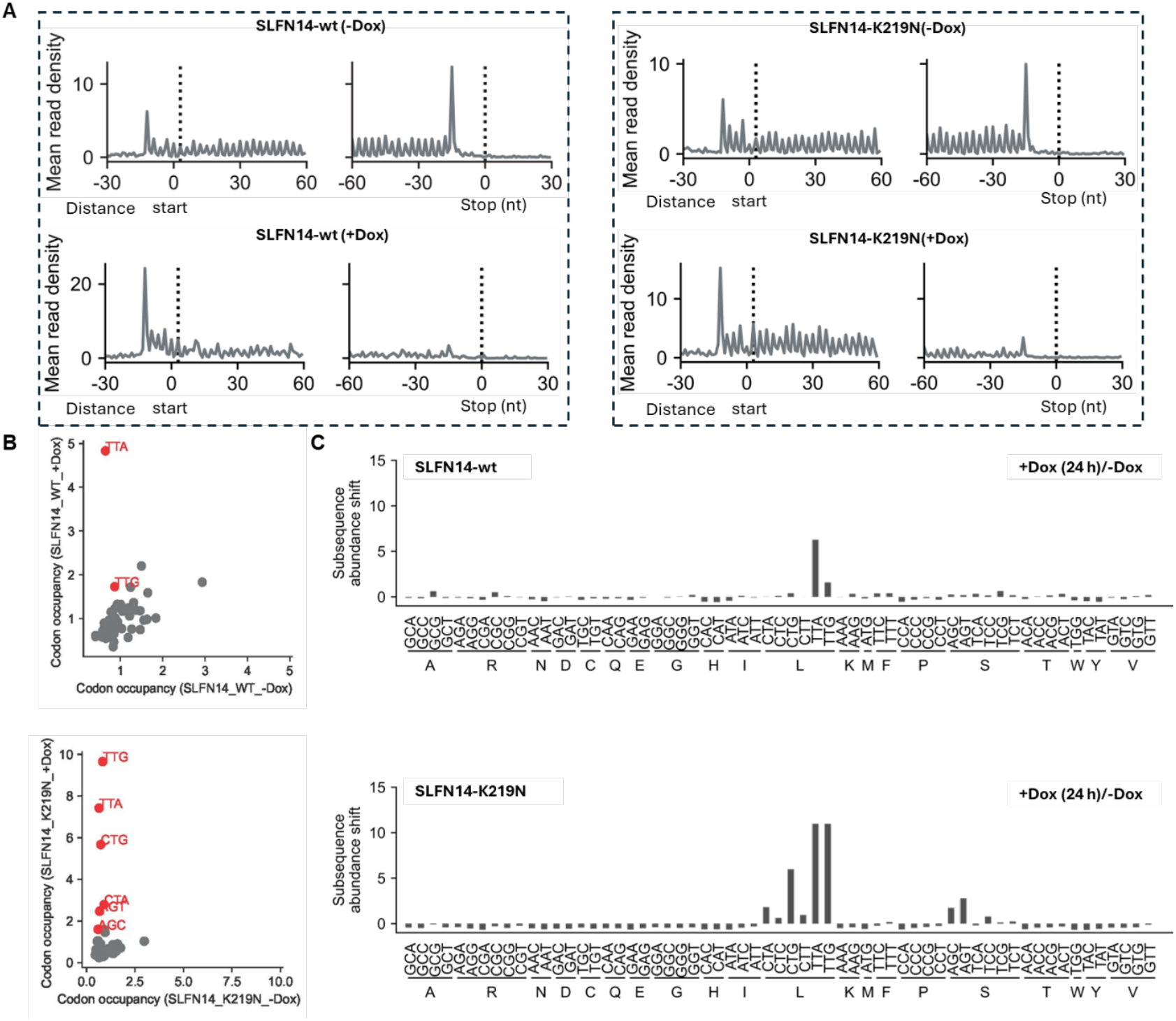
SLFN14 causes ribosomal stalling at codons decoded by type II tRNAs, an effect exacerbated by IT-linked mutations. Dox-inducible 293T cell lines expressing WT or K219N SLFN14 were left untreated or induced with Dox for 24 h before performing Ribo-seq. Global codon occupancy across all cellular transcripts was analyzed. **(A)** Aggregation plots around start and stop codons. All the samples showed clear 3-nt periodicity. **(B)** Codon occupancy analysis in Dox-induced cells relative to uninduced cells for each codon. Codons marked in red indicate those with large subsequence abundance shifts (> 1) under Dox treatment. (**C)** Subsequence abundance shifts of individual codons in Dox-induced cells relative to uninduced cells.

### IT-associated mutation causes increased cytotoxicity and eIF4E phosphorylation

While working with SLFN14 inducible cells, we frequently observed cell death after prolonged SLFN14 induction. To quantify this effect, we performed MTT assays at various time points after Dox induction of SLFN14. WT SLFN14 expression led to a gradual decrease in cell viability at 12 and 24 h post-induction, whereas the K219N variant caused an earlier reduction detectable at 6 h and a significantly greater decline at 24 h, indicating that the K219N mutation exacerbates SLFN14-induced cytotoxicity (Fig. 5A). This is also evident from time-course analysis of protein expression by Western blot. WT SLFN14 levels steadily increased after Dox induction, while housekeeping proteins (HSP70, β-actin, and GAPDH) remained stable. In contrast, induction of K219N for 24 h resulted in reduced β-actin and GAPDH levels, reflecting cell loss (Fig. 5B).

**Figure 5.**
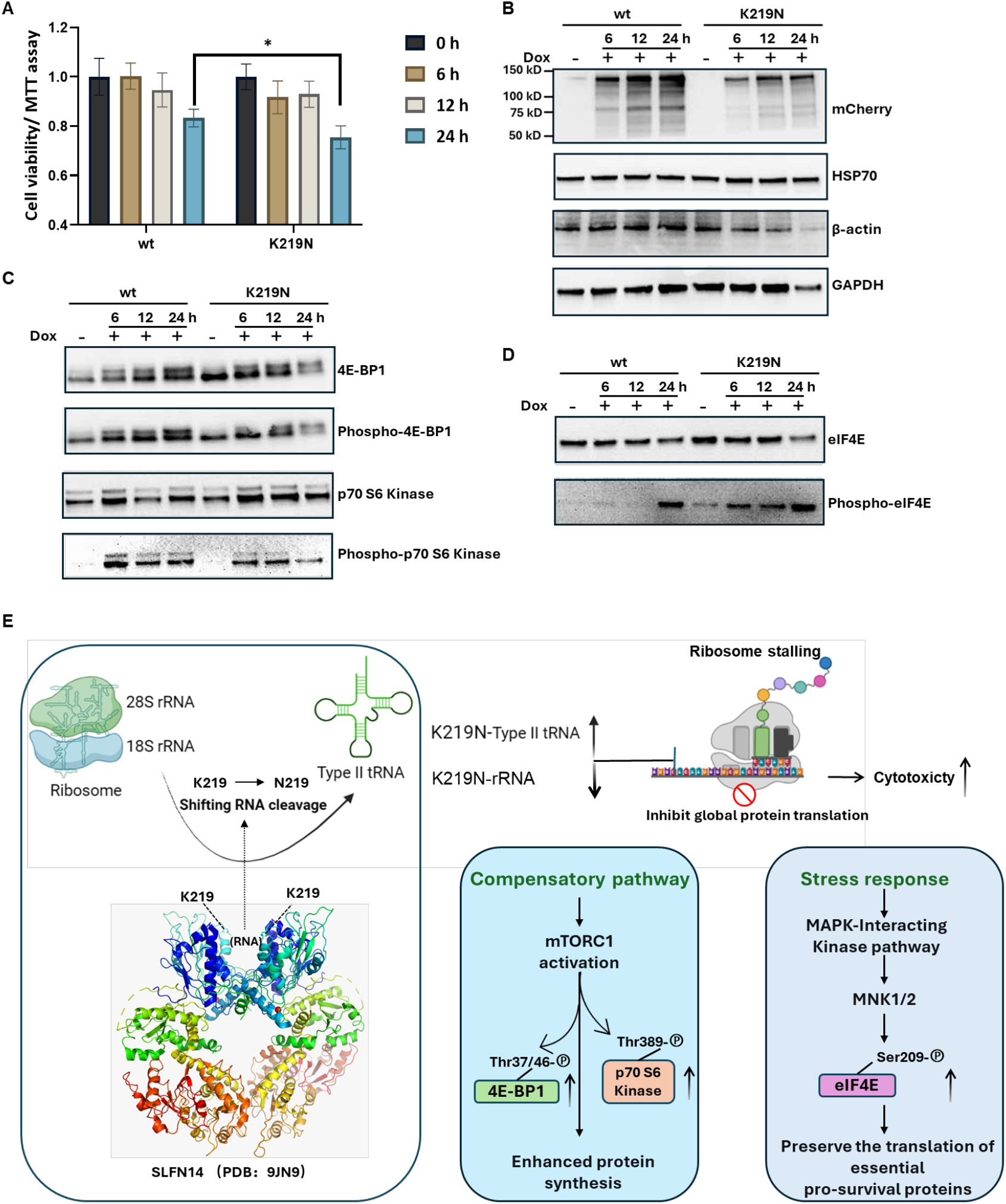
The K219N mutation in SLFN14 causes increased cytotoxicity and eIF4E phosphorylation. **(A)** 293T cells were induced to express SLFN14 for 6, 12, or 24 h, and cell viability was assessed using the MTT assay. **(B)** Western blot analysis of mCherry-SLFN14, HSP70, β-actin, and GAPDH expression. **(C)** Western blot analysis of mTORC1 pathway markers. **(D)** Western blot analysis of total eIF4E and phosphorylated eIF4E. **(E)** Schematic model illustrating how the IT-linked K219N mutation disrupts protein synthesis by shifting SLFN14 RNA cleavage activity toward type II tRNAs, leading to ribosome stalling, activation of compensatory stress pathways, and ultimately cell death.

A previous study suggested that SLFN14 activates the mTORC1 pathway as a compensatory response to translational dysregulation [3]. Activation of mTORC1 led to S6K and 4E-BP1 phosphorylation, and 4E-BP1 phosphorylation in turn relieves inhibition of eIF4E, a key cap-binding initiation factor. We found S6K and 4E-BP1 became phosphorylated within 6 hours of SLFN14 induction, indicating activation of the mTORC1 pathway (Fig. 5C). However, the magnitude of activation did not differ significantly between WT and K219N lines.

Beyond mTORC1 regulation, eIF4E activity is also regulated by phosphorylation to stimulate translation of some stress-responsive mRNAs [18]. Phosphorylated eIF4E was detected after 24 hours of WT SLFN14 induction, whereas in K219N cells, phosphorylation appeared earlier and progressively increased, suggesting that IT-linked SLFN14 mutations enhanced eIF4E phosphorylation as a stress response (Fig. 5D). These results suggest that, in addition to mTORC1 activation by SLFN14, MAPK pathway-driven eIF4E phosphorylation represents an additional compensatory stress response to the more severe translational arrest elicited by the K219N mutant (Fig. 5E).

### Overexpression of type II tRNAs counteracts SLFN14-mediated antiviral activity

To assess the contribution of tRNA depletion to SLFN14-mediated translational repression and antiviral activity, we co-expressed individual tRNAs in SLFN14-expressing cells and assessed VACV replication. VACV replication was quantified by both the percentage of cells expressing virus-encoded GFP and its MFI. Co-expression of any of the four tested tRNA modestly increased the percentage of GFP-positive cells in SLFN14-expressing cultures (Fig. 6A and C). Furthermore, two type II tRNA (tRNA-Leu^TAA^ and tRNA-Ser^GCT^) also significantly increased GFP intensity in cells expressing the K219N mutant (Fig. 6B and D), indicating higher levels of viral protein synthesis. These results indicate that depletion of type II tRNAs is particularly critical for K219N-mediated translational repression and antiviral activity.

**Figure 6.**
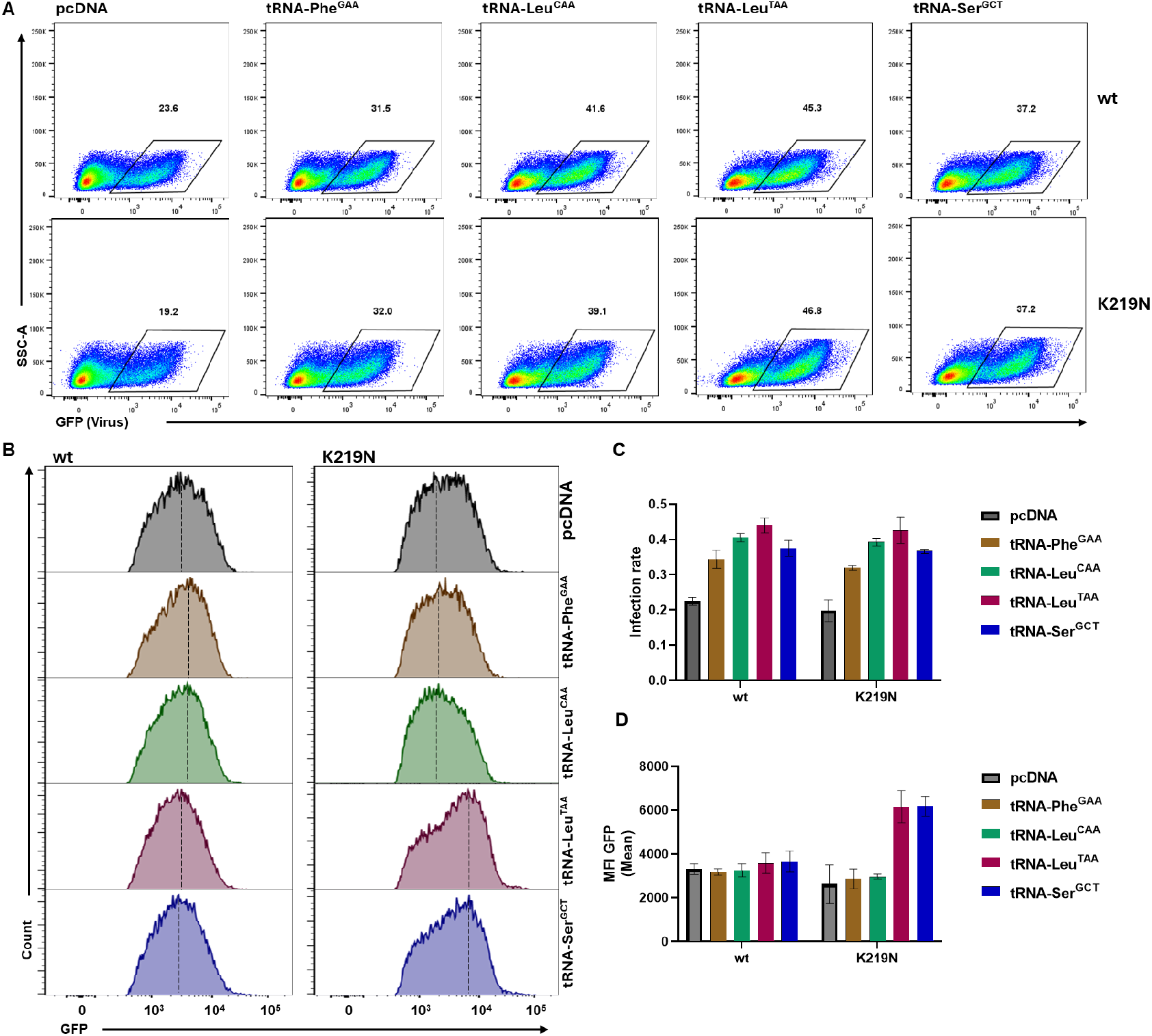
Overexpression of type II tRNAs counteracts the antiviral effect of SLFN14. HEK293T cells were transfected with mCherry-P2A-SLFN14 (WT or K219N) together with the indicated tRNA expression constructs for 8 h and then infected with VACV/GFP^+^ for 15 h. **(A)** Representative flow cytometry plots. **(B)** Histograms of GFP fluorescence intensity from the infected cell populations. **(C)** Quantification of the percentage of GFP+ cells shown in (A). **(D)** Quantification of GFP MFI in infected cell populations shown in (B).

## Discussion

Our study identifies impaired translation as a key pathogenic effect of IT-linked SLFN14 variants. Although these mutations were previously known to reduce SLFN14 protein expression, the underlying mechanism had remained unclear and was previously attributed to a dominant-negative effect on WT protein stability. Here, we demonstrate that reduced SLFN14 protein levels reflect enhanced translational repression by IT-linked variants, with relative potency in the order K218E ≈ K219N > R223W > WT > V220D. Apart from V220D, which was reported to behave differently from other IT-linked variants [5], the extent of translational inhibition by the variants roughly correlates with disease severity: patients carrying K218E or K219N mutations had similarly low platelet counts and severe bleeding [6], whereas those carrying R223W exhibit milder symptoms. This supports the idea that translational inhibition is the primary driver of platelet dysfunction in SLFN14-linked IT.

We further identify enhanced type II tRNA degradation as the molecular mechanism underlying the stronger translational block caused by IT-linked variants. SLFN14 is known to bind ribosomes and cleave rRNA, but there are conflicting reports on whether IT-linked mutations reduce or enhance rRNA degradation [3, 5]. Through detailed time-course studies, we found that the K219N variant decreases rRNA cleavage while enhancing tRNA degradation in cells. Notably, at earlier time points when translational inhibition became prominent, type II tRNAs were almost completely degraded by the K219N variant, whereas rRNA degradation remained minimal. The unbiased ribo-seq analysis revealed codon-specific ribosome stalling consistent with depletion of type II tRNAs. We propose that physiological SLFN14 activity involves cleavage of both rRNA and tRNA, but the K219N mutation impairs rRNA cleavage to a greater extent than tRNA cleavage, thereby shifting its activity towards tRNA degradation.

Low-level tRNA cleavage by WT SLFN14 in MK or platelets may be tolerated without impairing function, but IT-linked mutations increase tRNA degradation to levels that induce ribosomal stalling, stress responses, and ultimately cell death. Consistent with a previous report [3], we found that SLFN14 activated the mTORC1 pathway. However, while K219N did not further enhance mTORC1 activation, it enhanced eIF4E phosphorylation, known to promote translation of growth-promoting and stress-responsive mRNAs [18], suggesting an adaptive attempt to compensate for global translational repression.

Our findings also suggest that tRNA degradation is the primary antiviral mechanism of SLFN14. Although SLFN14 has been reported to cleave viral genomic RNAs in vitro [11, 12], we show that it can also effectively inhibit cytoplasmic DNA viruses. The enhanced antiviral activity of IT-linked variants underscores a direct correlation between tRNA cleavage, translational arrest, and viral restriction. By modulating codon-dependent translation through targeted depletion of type II tRNAs, interferon-induced SLFN14 may provide broad-spectrum antiviral activity, since all viruses depend on the host translational machinery.

The preferential targeting of type II tRNAs by SLFN14 suggests that this is a conserved regulatory strategy within the SLFN family. Although the RNase domains of SLFN11 and SLFN12 display broad RNA substrate specificity in vitro, the full-length proteins selectively target type II tRNAs, particularly tRNA-Leu^TAA^, in cells [19-21]. SLFN14 likewise has broad RNA substrate specificity in vitro and is known to target rRNA in cells, but here we show that its major cellular effect is the preferential depletion of type II tRNAs and ribosomal stalling at the corresponding codons, indicating that type II tRNAs are a common target for SLFN proteins.

SLFN11 is activated in response to DNA damage [20], whereas SLFN12 can be activated by small-molecule “molecular glues” [19]. In contrast, SLFN14 does not appear to require any activation signal beyond transcriptional induction.

The discovery that pathogenic SLFN14 variants inhibit global protein synthesis through selective tRNA depletion raises the possibility that modulation of tRNA homeostasis could mitigate translational arrest. For example, overexpression of type II tRNAs may represent a targeted therapeutic approach. Consistent with this idea, overexpression of type II tRNAs counteracts SLFN14 antiviral activity. Future studies using patient-derived megakaryocytes will be critical to test these strategies and to explore whether translation-modulating therapies can restore platelet production in SLFN14-associated thrombocytopenia.

## Materials and Methods

### Reagents

The antibodies against the following proteins were used in the immunoblotting: mCherry (Bio-techne, NBP2-25157), HSP70 (Santa Cruz, sc-137210), β-actin (Sigma, A2228), GAPDH (Cell Signaling Technology, #2118), 4E-BP1 (Cell Signaling Technology, #9452), Phospho-4E-BP1(Thr37/46) (Cell Signaling Technology, #2855), eIF4E (Cell Signaling Technology, #9742), Phospho-eIF4E (Ser209) (Cell Signaling Technology, #9741), p70 S6 Kinase (Cell Signaling Technology, #9202), Phospho-p70 S6 Kinase (Cell Signaling Technology, #9205). Additional reagents included high range RNA ladder (ThermoFisher, SM1821) and the Low Range ssRNA Ladder (NEB, N0364S).

### Cells and Viruses

Human embryonic kidney (HEK) 293T cells (ATCC CRL-3216) were cultured in Dulbecco’s Modified Eagle Medium (DMEM) supplemented with 10% fetal bovine serum (FBS) at 37ºC with 5.0% CO_2_. WT vaccinia virus (VACV) strain Western Reserve (WR) and the GFP-expressing VACV K1/C7 deletion mutant (vK1^-^C7^-^/GFP^+^) were described previously (*19-21*).

### Plasmid construction

Plasmids were constructed using PCR products or synthesized DNA fragments using the NEBuilder HiFi DNA Assembly Kit (NEB). pcDNA6.2/mCherry-p2A-hSLFN14 was constructed with DNA fragments synthesized by Twist Bioscience. Point mutations (K218E, K219N, V220D, R223W) were introduced using PCR products from synthesized oligos containing the desired mutations. mCherry-tagged SLFN14 WT and K219N sequences were PCR-amplified and cloned into a modified piggyBac vector with a Dox-inducible promoter and the reverse tetracycline transactivator [22] (Gift of Satoshi Narumi). Plasmids expressing tRNA-Leu^CAA^ and tRNA-Phe^GAA^ were described previously [23]. Plasmids expressing tRNA-Leu^TAA^ and tRNA-Ser^GCT^ were assembled using synthesized DNA fragments from Integrated DNA Technologies (IDT). All constructs were verified by whole-plasmid sequencing (Plasmidsaurus). Primer sequences are listed in Supplemental Table 1.

### Generation of Inducible SLFN14 Cell Lines

293T parental cells were transfected with 500 ng of the piggyBac vector encoding SLFN14 and 200 ng transposase expression plasmid (System Biosciences) in 6-well plates. After 72 h, cells were replated and selected with blasticidin (15 μg/mL) for 3 days, followed by single-cell sorting into 96-well plates. After 2 weeks, clones were screened for doxycycline (2 μg/mL) inducibility by mCherry expression. Positive clones were expanded, and SLFN14 expression was validated by immunoblotting with anti-mCherry antibody.

### Antiviral Assays with SLFN14 Variant

For transient-expression assays, HEK293T cells were transfected with plasmids expressing WT or mutant mCherry-SLFN14. After 24 h, cells were infected with vK1^-^C7^-^/GFP^+^ VACV for 15 h.

GFP expression was analyzed by fluorescence microscopy and quantified by flow cytometry, as described [24].

### Antiviral Assays in Inducible Cell Lines

WT or K219N SLFN14-inducible 293T cells were seeded in 24-well plates and left untreated or induced with doxycycline (2 μg/mL) for 24 h. Cells were infected with VACV (MOI 0.5) for 2 h at room temperature. After adsorption, cells were washed twice with PBS and incubated at 37°C. After 0, 24, and 48 h, samples were collected, and viral titers were determined by plaque assay on CV-1 cells. In some experiments, cells were also analyzed by flow cytometry.

### Assay of Nascent Protein Synthesis

Nascent protein synthesis was measured using the Click-iT Plus OPP Alexa Fluor 488 Protein Synthesis Assay Kit (Thermo Fisher Scientific). Cells were pulsed with O-propargyl-puromycin (OPP) for 30 min, followed by the Click-iT reaction and analysis by flow cytometry (BD LSR-II).

### Viral Rescue Assay with Co-transfection SLFN14 and Exogenous tRNA

HEK 293T cells were co-transfected with plasmids expressing SLFN14 mCherry fusion plasmids and a plasmid expressing tRNA-Phe^GAA^, tRNA-Leu^CAA^, tRNA-Leu^TAA^, or tRNA-Ser^GCT^ (pcDNA3.1 as control) for 12 h. Infection with VACV/GFP+ was conducted for 16 h and infection rates were determined with flow cytometry.

### RNA Analysis: rRNA and tRNA

Total RNA was extracted using TRIzol reagent. For rRNA analysis, 0.5 μg of RNA was resolved on a 2% agarose gel. For tRNA analysis, 2.5 μg of RNA was separated by 10% denaturing urea-PAGE and visualized with SYBR Gold staining.

### MTT assay

Cell viability of SLFN14 inducible 293T cells was tested using the MTT assay. Briefly, cells were seeded in 96-well plates and left untreated or treated with doxycycline (2 μg/mL) for indicated times (6, 12 or 24 h). 10 µL of MTT reagent (Invitrogen, V13154) was added to each well and incubated for 4 h at 37 °C. The reaction was terminated by adding 100 μL of DMSO, followed by a 10 min incubation at 37 °C. Absorbance was measured at 540 nm using a microplate reader (BioTek).

### Immunoblotting

Cell lysates were separated on 4-12% SDS-PAGE gradient gels (Bio-Rad) and transferred to nitrocellulose membranes (Thermo Fisher Scientific). Membranes were blocked with TBS containing 5% nonfat dry milk and 0.1% Tween-20 for 1 h at room temperature, incubated overnight at 4 °C with primary antibodies, and then probed with HRP-conjugated secondary antibodies. Bands were visualized by enhanced chemiluminescence (ECL-Plus, GE Healthcare).

### Northern Blotting

tRNAs were analyzed as described previously (*23*). Briefly, RNA samples were resolved on 15% TBE-urea gels, transferred to positively charged nylon membranes, and cross-linked by UV. DIG-labeled oligonucleotide probes specific for individual tRNAs (Supplemental Table 1) were hybridized to the membranes. Probes were generated using the DIG Oligonucleotide 3’-End Labeling Kit (Roche). Detection was performed using CDP-Star substrate (Roche) and chemiluminescence.

### Ribosome Profiling (Ribo-seq)

WT or K219N SLFN14-inducible 293T cells were seeded in 6-well plates and left untreated or induced with doxycycline (2 μg/mL) for 24 h. The 6-well plates were pre-coated with poly-D-lysine (Gibco) for 5 min at room temperature to enhance the cell attachment. Following treatment, cells were washed once with ice-cold PBS and lysed in polysome lysis buffer (10 mM HEPES, pH 7.4, 5 mM MgCl2, 100 mM KCl, 1% Triton X-100, 100 μg/mL cycloheximide).

Lysates were clarified by centrifugation at maximum speed (≥ 13,000 × g) for 15 min at 4 °C, and the supernatant was collected for downstream ribo-seq analysis. Ribo-seq libraries were constructed by modifying the Ezra-seq method, as previously described [25]. In brief, the whole cell lysates were digested with E. coli RNase I (Ambion, 750 U per 100 A260 units) by agitation at 4°C for 1 h. RNAs were extracted using Trizol LS reagent (Invitrogen) followed by isopropanol precipitation. The ribosome-protected mRNA fragments (RPFs) were separated on a 15% polyacrylamide TBE-urea gel (Invitrogen) and visualized using SYBR Gold (Invitrogen).

Selected regions in the gel corresponding to 25-35 nt were excised and dissolved by soaking in 400 μl RNA elution buffer (300 mM NaOAc pH 5.2, 1 mM EDTA, 0.1 U/μl SUPERase·In (Invitrogen)) for 10 min at 70 °C. The gel debris was removed using a Spin-X column (Corning), followed by isopropanol precipitation. 4 μl RNAs (10∼200 ng) were mixed with 1 nmol ATP, 1 μl T4 PNK (NEB), 20 U SUPERase·In, 1 μl Poly(A) Polymerase (NEB), and 1 μl homemade Ezra enzyme in 1 × T4 PNK buffer and incubated at 37 °C for 30 min followed by 70 °C for 10 min. Then ligation was performed for 90 min at 25 °C by mixing with a 10 μl reaction mixture (1 pmol biotinylated 5’ end adaptor, 1 × T4 Rnl2 reaction buffer, 20 U SUPERase·In, 12.5% PEG8000 and 200 U T4 RNA ligase 2 truncated KQ (NEB)). The ligated RNA samples were pulled down after incubation with 20 uL pre-washed streptavidin magnetic beads (NEB) at room temperature for 10 min. After washing once with 100 uL 2 × SSC (Invitrogen), beads were resuspended in 12 μl nuclease-free water and mixed with 8 μl cDNA synthesis mixture (1 poml RT primer, 1 × first strand buffer, 100 nmol DTT, 1 nmol dNTP, 20 U SUPERase·In and 100 ng m-MLV) followed by incubation at 50 °C for 20 min. After washing once with 2 × SSC, beads were resuspended in 20 μl nuclease-free water and incubated at 95 °C for 2 min, then immediately placed on the ice for 1 min. After placing on magnet stand for 1 min, the supernatant cDNA was amplified by PCR using barcoded sequencing primers. PCR was performed by mixing 1 × HF buffer, 0.5 mM dNTP, 0.25 μM PCR primers and 0.025 U Phusion polymerase. PCR was carried out under the following conditions: 98 °C, 30 s; (98 °C, 5 s; 67 °C, 15 s; 72 °C, 20 s) for 13 cycles; 72 °C, 2 min. PCR products were separated on an 8% polyacrylamide TBE gel (Invitrogen). DNA products with the expected size 170 bp were excised and recovered from DNA elution buffer (300 mM NaCl, 1 mM EDTA). After quantification by Qubit 4 Fluorometer (Invitrogen), equal amounts of barcoded samples were pooled and sequenced using MiniSeq (Illumina). The oligonucleotide sequences are listed in Supplemental Table 1.

### Ribo-seq analysis

To align sequencing reads, the 5’ and 3’ adapters of the reads were trimmed by Cutadapt (version 2.8) [26]. The trimmed reads with length shorter than 15 nucleotides were excluded from the analysis. To keep accurate reading frame of Ribo-seq, low-quality bases at both ends of the reads were not subject to clip. The trimmed reads were first aligned to rRNAs using Bowtie (version 1.2.3) [27]. The rRNA sequences were downloaded from the nucleotide database of NCBI and RNAcentral. The reads unaligned to rRNAs were then mapped to the custom human transcriptome using STAR (version 2.7.10a) [28]. To avoid ambiguity, reads mapped to multiple positions or with >2 mismatches were disregarded for further analysis. The custom transcriptome was generated based on the reference genome and annotations obtained from Ensembl using human release 109 (GRCh38.p14). Protein coding genes were extracted, and a single transcript was selected for each gene on the following procedure. For each gene, the transcript with the highest APPRIS score was initially selected. If the selected transcripts have equal APPRIS scores, the transcript with the longest coding sequence (CDS) was included in the custom transcriptome. Mapping to ribosomal P sites was performed by shifting the read position from the 5’ ends to the position by 12 nt.

For codon occupancy analysis, ribosome density at each codon was calculated by normalizing the read count with the average read counts per codon of the transcript, and then was averaged across the transcriptome. The subsequence abundance shift plots were created following diricore analysis [29].

### Quantification and Statistical Analysis

Data were analyzed using GraphPad Prism 9. Statistical methods, including one-way ANOVA, are specified in the figure legends.

## Data availability

All data are available in the main text or the supplementary materials. Ribo-seq data are deposited in the NCBI GEO under accession number GSE310203 (reviewer token:mlqjwymyhfynval). The code used for the Ribo-seq analysis is deposited to GitHub (https://github.com/usa0ri/SLFN14)

## Competing interests

The authors declare no competing financial interests.

### Author contributions

Conceptualization: YX,

CD Methodology: CD,FZ, XL, SU

Investigation: CD,FZ, XL, SU

Visualization: CD, XL, SU

Funding acquisition: YX

Project administration: YX, SQ

Supervision: YX, SQ

Writing – original draft: YX

Writing – review & editing: CD, XL, FZ, SU,SQ,YX

## References

1. Jo U, Pommier Y. Structural, molecular, and functional insights into Schlafen proteins. Exp Mol Med. 2022;54(6):730–8. Epub 20220629. doi: 10.1038/s12276-022-00794-0. PubMed PMID: 35768579; PubMed Central PMCID: PMCPMC9256597.

2. Kim ET, Weitzman MD. Schlafens Can Put Viruses to Sleep. Viruses. 2022;14(2). Epub 20220221. doi: 10.3390/v14020442. PubMed PMID: 35216035; PubMed Central PMCID: PMCPMC8875196.

3. Ver Donck F, Ramaekers K, Thys C, Van Laer C, Peerlinck K, Van Geet C, et al. Ribosome dysfunction underlies SLFN14-related thrombocytopenia. Blood. 2023;141(18):2261–74. doi: 10.1182/blood.2022017712. PubMed PMID: 36790527; PubMed Central PMCID: PMCPMC10646786.

4. Stapley RJ, Smith CW, Haining EJ, Bacon A, Lax S, Pisareva VP, et al. Heterozygous mutation SLFN14 K208N in mice mediates species-specific differences in platelet and erythroid lineage commitment. Blood Adv. 2021;5(2):377–90. doi: 10.1182/bloodadvances.2020002404. PubMed PMID: 33496736; PubMed Central PMCID: PMCPMC7839357.

5. Fletcher SJ, Pisareva VP, Khan AO, Tcherepanov A, Morgan NV, Pisarev AV. Role of the novel endoribonuclease SLFN14 and its disease-causing mutations in ribosomal degradation. RNA. 2018;24(7):939–49. Epub 20180420. doi: 10.1261/rna.066415.118. PubMed PMID: 29678925; PubMed Central PMCID: PMCPMC6004054.

6. Fletcher SJ, Johnson B, Lowe GC, Bem D, Drake S, Lordkipanidze M, et al. SLFN14 mutations underlie thrombocytopenia with excessive bleeding and platelet secretion defects. J Clin Invest. 2015;125(9):3600–5. Epub 20150817. doi: 10.1172/JCI80347. PubMed PMID: 26280575; PubMed Central PMCID: PMCPMC4588283.

7. Marconi C, Di Buduo CA, Barozzi S, Palombo F, Pardini S, Zaninetti C, et al. SLFN14-related thrombocytopenia: identification within a large series of patients with inherited thrombocytopenia. Thromb Haemost. 2016;115(5):1076–9. Epub 20160114. doi: 10.1160/TH15-11-0884. PubMed PMID: 26769223.

8. Polokhov D, Fedorova D, Ignatova A, Ponomarenko E, Rashevskaya E, Martyanov A, et al. Novel SLFN14 mutation associated with macrothrombocytopenia in a patient with severe haemorrhagic syndrome. Orphanet J Rare Dis. 2023;18(1):74. Epub 20230411. doi: 10.1186/s13023-023-02675-9. PubMed PMID: 37041648; PubMed Central PMCID: PMCPMC10091655.

9. Seong RK, Seo SW, Kim JA, Fletcher SJ, Morgan NV, Kumar M, et al. Schlafen 14 (SLFN14) is a novel antiviral factor involved in the control of viral replication. Immunobiology. 2017;222(11):979–88. Epub 20170711. doi: 10.1016/j.imbio.2017.07.002. PubMed PMID: 28734654; PubMed Central PMCID: PMCPMC5990420.

10. Valenzuela C, Saucedo S, Llano M. Schlafen14 Impairs HIV-1 Expression in a Codon Usage-Dependent Manner. Viruses. 2024;16(4). Epub 20240325. doi: 10.3390/v16040502. PubMed PMID: 38675845; PubMed Central PMCID: PMCPMC11054720.

11. Li M, Sun D, Hao W, Fu H, Zhang Y, Li Z, et al. Human Schlafen 14 Cleavage of Short Double-Stranded RNAs Underpins its Antiviral Activity. Adv Sci (Weinh). 2025:e01727. Epub 20250711. doi: 10.1002/advs.202501727. PubMed PMID: 40642785.

12. Luo M, Jia X, Wang ZW, Yang JY, Wang W, Chen J, et al. Structural and functional characterization of human SLFN14. Nucleic Acids Res. 2025;53(10). doi: 10.1093/nar/gkaf484. PubMed PMID: 40464691; PubMed Central PMCID: PMCPMC12135189.

13. Van Riper J, Martinez-Claros AO, Wang L, Schneiderman HE, Maheshwari S, Pillon MC. CryoEM structure of the SLFN14 endoribonuclease reveals insight into RNA binding and cleavage. Nature communications. 2025;16(1):5848. Epub 20250701. doi: 10.1038/s41467-025-61091-8. PubMed PMID: 40592880; PubMed Central PMCID: PMCPMC12215978.

14. Yang JY, Deng XY, Li YS, Ma XC, Feng JX, Yu B, et al. Structure of Schlafen13 reveals a new class of tRNA/rRNA-targeting RNase engaged in translational control. Nature communications. 2018;9(1):1165. Epub 20180321. doi: 10.1038/s41467-018-03544-x. PubMed PMID: 29563550; PubMed Central PMCID: PMCPMC5862951.

15. Garvie CW, Wu X, Papanastasiou M, Lee S, Fuller J, Schnitzler GR, et al. Structure of PDE3A-SLFN12 complex reveals requirements for activation of SLFN12 RNase. Nature communications. 2021;12(1):4375. Epub 20210716. doi: 10.1038/s41467-021-24495-w. PubMed PMID: 34272366; PubMed Central PMCID: PMCPMC8285493.

16. Metzner FJ, Huber E, Hopfner KP, Lammens K. Structural and biochemical characterization of human Schlafen 5. Nucleic Acids Res. 2022;50(2):1147–61. doi: 10.1093/nar/gkab1278. PubMed PMID: 35037067; PubMed Central PMCID: PMCPMC8789055.

17. Metzner FJ, Wenzl SJ, Kugler M, Krebs S, Hopfner KP, Lammens K. Mechanistic understanding of human SLFN11. Nature communications. 2022;13(1):5464. Epub 20220917. doi: 10.1038/s41467-022-33123-0. PubMed PMID: 36115853; PubMed Central PMCID: PMCPMC9482658.

18. Yang X, Zhong W, Cao R. Phosphorylation of the mRNA cap-binding protein eIF4E and cancer. Cell Signal. 2020;73:109689. Epub 20200611. doi: 10.1016/j.cellsig.2020.109689. PubMed PMID: 32535199; PubMed Central PMCID: PMCPMC8049097.

19. Lee S, Hoyt S, Wu X, Garvie C, McGaunn J, Shekhar M, et al. Velcrin-induced selective cleavage of tRNA(Leu)(TAA) by SLFN12 causes cancer cell death. Nat Chem Biol. 2023;19(3):301–10. Epub 20221027. doi: 10.1038/s41589-022-01170-9. PubMed PMID: 36302897.

20. Boon NJ, Oliveira RA, Körner P-R, Kochavi A, Mertens S, Malka Y, et al. DNA damage induces p53-independent apoptosis through ribosome stalling. Science. 2024;384(6697):785–92. doi: doi:10.1126/science.adh7950.

21. Li M, Kao E, Malone D, Gao X, Wang JYJ, David M. DNA damage-induced cell death relies on SLFN11-dependent cleavage of distinct type II tRNAs. Nat Struct Mol Biol. 2018;25(11):1047–58. Epub 2018/10/31. doi: 10.1038/s41594-018-0142-5. PubMed PMID: 30374083; PubMed Central PMCID: PMCPMC6579113.

22. Shima H, Koehler K, Nomura Y, Sugimoto K, Satoh A, Ogata T, et al. Two patients with MIRAGE syndrome lacking haematological features: role of somatic second-site reversion SAMD9 mutations. J Med Genet. 2018;55(2):81–5. Epub 20171124. doi: 10.1136/jmedgenet-2017-105020. PubMed PMID: 29175836.

23. Zhang F, Ji Q, Chaturvedi J, Morales M, Mao Y, Meng X, et al. Human SAMD9 is a poxvirus-activatable anticodon nuclease inhibiting codon-specific protein synthesis. Science Advances. 2023;9(23):eadh8502. doi: doi:10.1126/sciadv.adh8502.

24. Morales M, Zhang F, Xiang Y. Investigating the Impact of Host Factors on Vaccinia Virus Infection Through Single-Cell Analysis via Flow Cytometry. Methods Mol Biol. 2025;2860:219–27. doi: 10.1007/978-1-0716-4160-6_14. PubMed PMID: 39621270.

25. Mao Y, Jia L, Dong L, Shu XE, Qian SB. Start codon-associated ribosomal frameshifting mediates nutrient stress adaptation. Nat Struct Mol Biol. 2023;30(11):1816–25. Epub 20231113. doi: 10.1038/s41594-023-01119-z. PubMed PMID: 37957305.

26. Martin M. Cutadapt removes adapter sequences from high-throughput sequencing reads. 2011. 2011;17(1):3. Epub 2011-08-02. doi: 10.14806/ej.17.1.200.

27. Langmead B, Trapnell C, Pop M, Salzberg SL. Ultrafast and memory-efficient alignment of short DNA sequences to the human genome. Genome Biology. 2009;10(3):R25. doi: 10.1186/gb-2009-10-3-r25.

28. Dobin A, Davis CA, Schlesinger F, Drenkow J, Zaleski C, Jha S, et al. STAR: ultrafast universal RNA-seq aligner. Bioinformatics. 2013;29(1):15–21. Epub 20121025. doi: 10.1093/bioinformatics/bts635. PubMed PMID: 23104886; PubMed Central PMCID: PMCPMC3530905.

29. Loayza-Puch F, Rooijers K, Buil LC, Zijlstra J, Oude Vrielink JF, Lopes R, et al. Tumour-specific proline vulnerability uncovered by differential ribosome codon reading. Nature. 2016;530(7591):490–4. Epub 20160215. doi: 10.1038/nature16982. PubMed PMID: 26878238.

